# The visual coupling between neighbors explains ‘flocking’ in human crowds

**DOI:** 10.1101/2021.09.04.459001

**Authors:** Gregory C. Dachner, Trenton D. Wirth, Emily Richmond, William H. Warren

## Abstract

Patterns of collective motion or ‘flocking’ in birds, fish schools, and human crowds are believed to emerge from local interactions between individuals. Most models of collective motion attribute these interactions to hypothetical rules or forces, often inspired by physical systems, and described from an overhead view. We develop a *visual model* of human flocking from an embedded view, based on optical variables that actually govern pedestrian interactions. Specifically, people control their walking speed and direction by canceling the average optical expansion and angular velocity of their neighbors, weighted by visual occlusion. We test the model by simulating data from experiments with virtual crowds and real human ‘swarms’. The visual model outperforms our previous overhead model and explains basic properties of physics-inspired models: ‘repulsion’ forces reduce to canceling optical expansion, ‘attraction’ forces to canceling optical contraction, and ‘alignment’ to canceling the combination of expansion/contraction and angular velocity. Critically, the neighborhood of interaction follows from Euclid’s Law of perspective and the geometry of occlusion. We conclude that the local interactions underlying human flocking are a natural consequence of the laws of optics. Similar principles may apply to collective motion in other species.

## Introduction

Human crowds exhibit patterns of collective motion in many public settings, from train stations and shopping plazas to – sometimes catastrophically – mass events (Helbing, Buzna, Johansson, & Werner, 2005; Ngai, Burkle, Hsu, & Hsu, 2009). Similar patterns of coordinated motion are observed in bird flocks, fish schools, and animal herds, suggesting that diverse systems may obey common principles of self-organization (Couzin & Krause, 2003; Sumpter, 2010). It is generally believed that these global ‘flocking’ patterns emerge from local interactions between individuals (Couzin & Krause, 2003; Sumpter, 2010; Vicsek & Zafeiris, 2012). The crux of the problem thus lies in understanding the nature of the local interactions.

Most models of collective motion ascribe these local interactions to hypothetical rules or forces, often inspired by physical systems (Giardina, 2008; Schellinck & White, 2011). Such descriptive models – including our own (Rio, Dachner, & Warren, 2018) – assume an overhead view of the positions and velocities of all individuals in space. But humans and animals are immersed in a group and coupled by sensory information. Here we develop a visual model of collective motion that explains local interactions in terms of optical variables, from the viewpoint of an embedded agent. Not only does this explanatory model outperform our previous descriptive model, but basic properties of interaction turn out to be natural consequences of the laws of optics.

Understanding the local interactions involves, first, identifying the *rules of engagement* that govern how an individual responds to a neighbor, and second, characterizing the *neighborhood of interaction* over which the influences of multiple neighbors are combined. Classical “zonal” models (Couzin, Krause, James, Ruxton, & Franks, 2002; Huth & Wissel, 1992; Reynolds, 1987) assume three local rules or forces in concentric zones: (i) *repulsion* from neighbors in a near zone to avoid collisions, (ii) *alignment* with the velocity of neighbors in an intermediate zone to generate common motion, and (iii) *attraction* to neighbors in a far zone to ensure group cohesion. Neighbor influences are combined by averaging within a zone (Couzin et al., 2002; Huth & Wissel, 1992; Reynolds, 1987), often weighted by neighbor distance (Cucker & Smale, 2007; Grégoire, Chaté, & Tu, 2003; Mogilner, Edelstein-Keshet, Bent, & Spiros, 2003), and obey the superposition principle, according to which the response to a group is the linear combination of independent responses to each neighbor. An alignment rule by itself is theoretically sufficient to generate collective motion (Leonard et al., 2012; Vicsek, Czirók, Ben-Jacob, Cohen, & Shochet, 1995), as is the combination of attraction and repulsion (Romanczuk, Couzin, & Schimansky-Geier, 2009). Among pedestrian models (Chraibi, Tordeux, Schadschneider, & Seyfried, 2018), the prominent Social Force model (Chen, Treiber, Kanagaraj, & Li, 2018; Helbing & Molnár, 1995; Hoogendoorn & Bovy, 2003) is predicated on attraction and repulsion forces and has convenient mathematical properties (Köster, Treml, & Gödel, 2013). The model successfully simulates key crowd scenarios (Boltes, Zhang, Tordeux, Schadschneider, & Seyfried, 2018; Chraibi et al., 2018) and can generate collective motion under certain conditions (Helbing, Farkas, & Vicsek, 2000; Helbing, Molnár, Farkas, & Bolay, 2001), but does not produce realistic human trajectories (Pelechano, Allbeck, & Badler, 2007) or generalize to novel situations without re-parameterization (Campanella, Hoogendoorn, & Daamen, 2009; Chen et al., 2018).

The strength of such physics-inspired models is that they capture generic properties of collective motion, but the metaphor does not transfer smoothly to biological agents (Schellinck & White, 2011; Weitz et al., 2012). Because different sets of rules can produce similar global patterns (Vicsek & Zafeiris, 2012; Weitz et al., 2012), some researchers have turned to a ‘bottom-up’ approach that deploys experimental studies of behavioral interactions to infer the local rules (Calovi et al., 2018; Gautrais et al., 2012; Katz, Tunstrøm, Ioannou, Huepe, & Couzin, 2011; Moussaid et al., 2009; Sumpter, Mann, & Perna, 2012; Warren & Fajen, 2008; Zienkiewicz, Barton, Porfiri, & Di Bernardo, 2015). We further argue that a successful bottom-up model must be grounded in the sensory information that actually governs these interactions.

Initially, limits on the field of view and sensory range were introduced to constrain models of local interactions (Huth & Wissel, 1992; Partridge & Pitcher, 1980; Pita, Collignon, Halloy, & Fernández-Juricic, 2016). An analysis of fish schools discovered that a neighborhood based on visibility accounted for individual responses better than zonal or topological neighborhoods (Strandburg-Peshkin et al., 2013). Local interactions also strongly depend on the optical information that controls locomotion (Gibson, 1979; Pepping & Grealy, 2007; Warren, 1998). This insight was taken up by ‘vision-based’ (Dutra, Marques, Cavalcante-Neto, Vidal, & Pettré, 2017; Ondrej, Pettré, Olivier, & Donikian, 2010) and ‘cognitive heuristic’ (Bailo, Carrillo, & Degond, 2018; Moussaïd, Helbing, & Theraulaz, 2011) models, which were based on properties such as time-to-closest-approach, distance-at-closest-approach, or an interaction force related to time-to-contact (Karamouzas, Skinner, & Guy, 2014); however, the underlying optical variables were not specified. Recently, a minimal theoretical approach has been developed (Bastien & Romanczuk, 2020), which starts with simple visual directions as input and studies the emergent behavior in agent-based simulations.

We take a complementary, experiment-driven approach called ‘behavioral dynamics’ (Warren, 2006; Warren & Fajen, 2008), in which control is based on higher-order optical variables that animals are known to exploit(Frost & Sun, 2004; Srinivasan, 1998; Warren, Kay, Zosh, Duchon, & Sahuc, 2001). To model collective motion from the bottom up, we began with experiments on following in pedestrian dyads (Dachner & Warren, 2014; Rio, Rhea, & Warren, 2014). The results revealed that humans obey an alignment rule: a follower tends to match the walking direction (*heading*) and speed of the leader. We then mapped the neighborhood of interaction by asking participants to walk with a crowd in virtual reality, which allowed us to precisely control the movements of their virtual neighbors. The results showed that pedestrians average the heading directions and speeds of neighbors within the field of view, while analysis of naturalistic data from a human ‘swarm’ found that neighbor influence decays exponentially with distance, going to zero at about 4-5m (Rio et al., 2018).

These findings led to Rio, Dachner & Warren’s (2018) descriptive *behavioral model* of collective motion (Figure 1a; see SI). In brief, steering (angular acceleration) is controlled by canceling the difference between one’s current heading and the weighted average of neighbor headings. Speed (linear acceleration) is similarly controlled by canceling the difference between current speed and the weighted average of neighbor speeds. The weights decay exponentially with neighbor distance within a 180° field of view, yielding a semi-circular ‘soft metric’ neighborhood that generates robust collective motion in simulation (Cucker & Smale, 2007; Warren & Dachner, 2018). The model was fit to our data on pedestrian dyads, with the decay rate taken from the human swarm, and successfully predicted individual trajectories in virtual crowd experiments and real crowd data.

**Figure 1.**
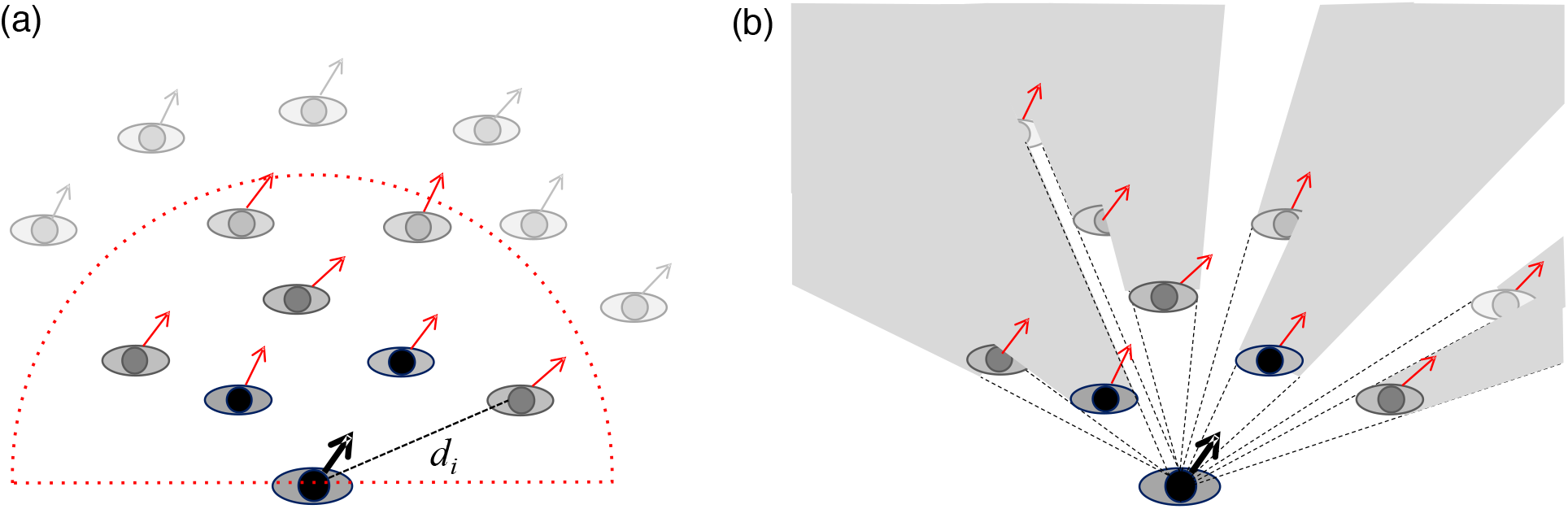
Overhead and embedded models of collective motion. (a) Overhead behavioral model: a pedestrian (bottom) matches the average heading direction of all neighbors in a 180° neighborhood. Neighbor weights (gray level) decay exponentially with distance *d_i_* and go to zero at 4-5m (dotted red curve). (b) Embedded visual model: a pedestrian (bottom) cancels the average angular velocity and optical expansion/contraction of all visible neighbors. Neighbor influence (gray level) decreases due to Euclid’s law of perspective and is proportional to visibility (shaded areas indicate occluded regions).

The behavioral model describes an empirical relation between physical variables – the velocity of an individual and the positions and velocities of neighbors. Like its predecessors, it assumes an overhead view, relies on metaphorical forces, and does not explain the local interactions. To address these issues, we developed a *visual model* predicated on the optical variables that are known to control pedestrian following (Figure 1b). This new model explains local interactions and the form of the neighborhood as resulting from the laws of optics and the geometry of visual occlusion.

## Results

### Behavioral results

#### The neighborhood of interaction is not fixed

According to the behavioral model, the neighborhood of interaction has a fixed radius, such that the influence of neighbors decays to zero at 4-5m (Rio et al., 2018). We tested this limit in a simple experiment by manipulating the distance to a virtual crowd and perturbing heading direction. To elicit collective-motion responses, participants (n=12) were instructed to “walk with the group of virtual humans” and “treat them as if they were real people.” A row of virtual humans (2, 4, or 8) was presented in a wireless head-mounted display, and its initial distance was manipulated (1.8, 3.0, 4.0, 6.0 or 8.0 m) (Figure 2a). On each trial, they appeared with their backs to the participant, began walking forward (1.0 m/s), and after 5s all turned by 10° (left or right) in the same direction and continued walking for another 7s. There was thus no occlusion. We used a head tracker to record the participant’s position and computed a time series of heading for each trial. The dependent measure was the participant’s *final heading*, the average heading direction during the last 2s of each trial (left/right turns were collapsed).

**Figure 2.**
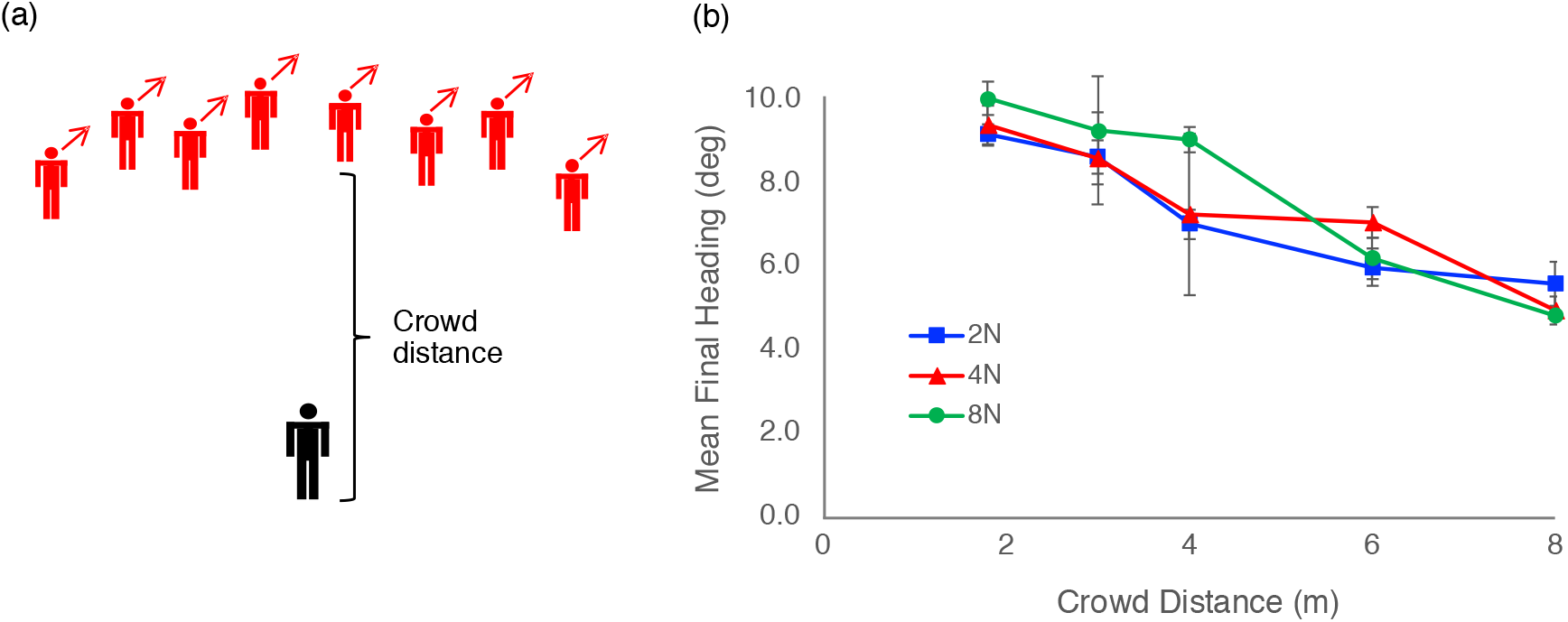
First experiment: Gradual decay to nearest neighbors. (a) Schematic of virtual crowd, illustrating a heading perturbation of all neighbors to the right. The distance between the participant (bottom) and the row of neighbors (top), and the number of neighbors, were manipulated. (b) Results: Mean final heading as a function of crowd distance. Curves represent the number of neighbors in the crowd; error bars represent ±SE.

To our surprise, we observed a very gradual decay in neighbor influence over a long distance (Figure 2b). The participant’s response decreased from a maximum final heading (*M*=9.55°) at 1.8m to just half that value (*M*=5.16°) at 8m, (*F* (4, 44)=14.93, *p*<0.001, 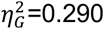). Linear extrapolation implies that the response would decay to zero around 15m (*y* = −0.722*x* + 10.8, *r* (14) *=* −0.95). There was no effect of crowd size on final heading (*F* (2, 22)=0.77, *p*=0.476, 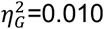) and no interaction (*F* (8,88)=0.83 *p*=0.575, 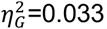), consistent with weighted averaging of neighbors.

The results clearly show that, contrary to a neighborhood with a fixed radius of 4-5m, pedestrians interact with neighbors at much greater distances – when they are not occluded. These findings imply that there may be two decay rates with distance: a gradual decay to the nearest neighbors, and a more rapid decay within a crowd. We tested this ‘double-decay’ hypothesis in the next experiment.

#### The double-decay hypothesis

To test two decay processes with distance, we asked each participant (n=10) to walk with a virtual crowd of 12 neighbors, positioned in three rows (Figure 3a). We replicated the decay to the nearest neighbors by manipulating the distance of the near row (2, 4, or 6 m from the participant), and probed the decay within the crowd by selectively perturbing the near, middle, or far row (always 2m apart). At the perturbation, all neighbors in one row turned by 10° in the same direction. Otherwise, the procedure was the same as before.

**Figure 3.**
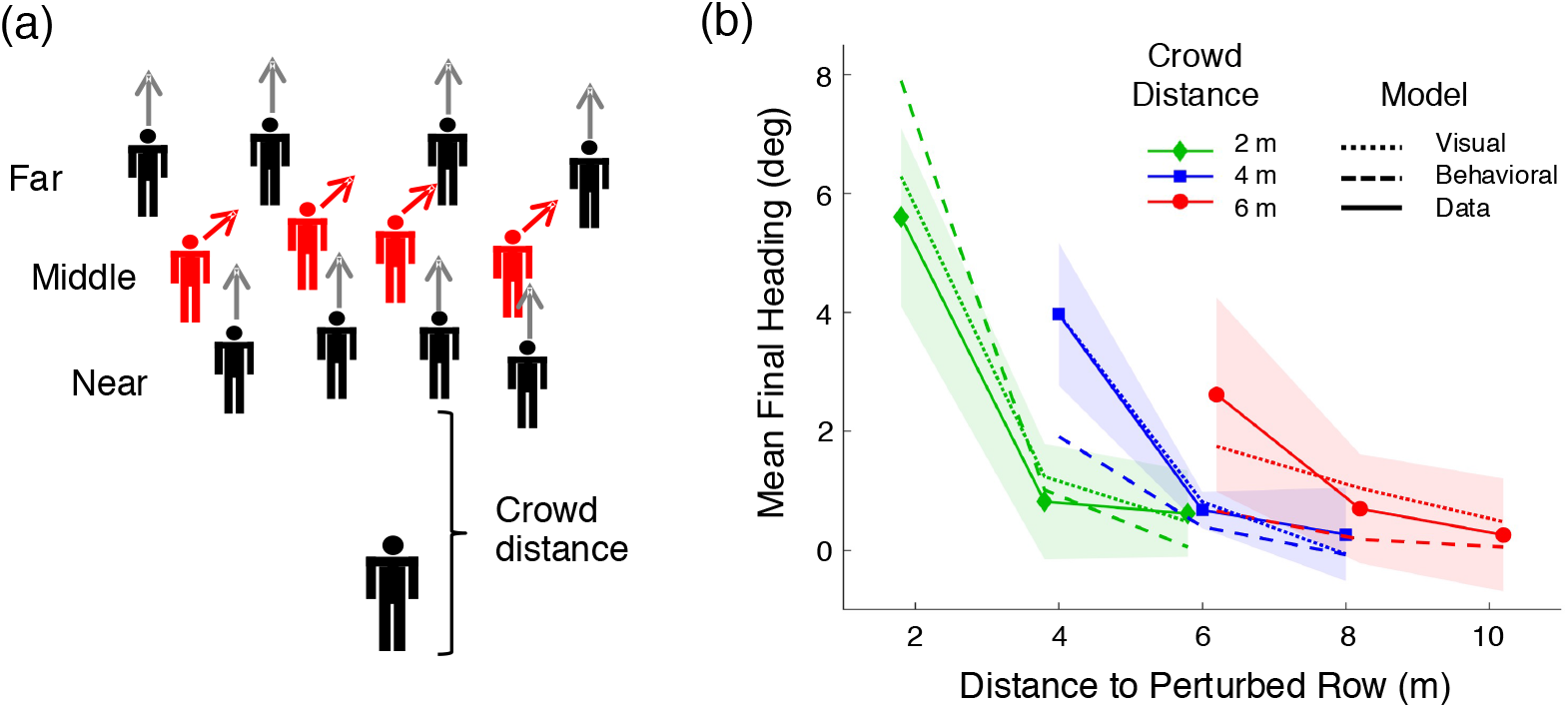
Second experiment: Test of the double-decay hypothesis. (a) Schematic of virtual crowd, illustrating a heading perturbation of one row to the right. The distance to the crowd (near row), and the perturbed row (near, middle, far), were manipulated. (b) Results: Mean final heading as a function of distance to the perturbed row, for each crowd distance (curves). Solid curves correspond to the human data, dotted curves to the visual model, and dashed curves to the behavioral model with the double-decay function. Shaded regions represent 95% confidence intervals for the human data; because the models are not intended to reproduce gait oscillations, their variable error is small and is not represented in the graph.

Figure 3b plots mean final heading as a function of distance to the perturbed row, where each curve represents a crowd distance (2, 4, or 6m). Two decay rates are immediately apparent. First, the heading response decreases with the overall distance of the crowd, (*F* (2,18)=8.59, *p*=0.002, 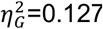); in particular, the response to perturbations of the near row (leftmost point of each curve) decays gradually with distance (simple effect test, *F* (2,18)=12.196, *p*<0.001), replicating the previous experiment. Linear extrapolation suggests that responses would go to zero around 10m (*y* = -.44x + 4.5, *r* (2) = -.97). Second, in each curve the response decreases rapidly within the crowd, (*F* (2,18)=35.64, *p*<0.001, 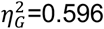), steeply from the near row to the middle row (Bonferroni-corrected *t* (9)=8.822, p<0.001) and the far row (*t* (9)=7.718, *p*<0.001), which approaches zero. This finding suggests that partial occlusion by near neighbors significantly weakened the influence of middle neighbors, and far neighbors were almost completely occluded. In a separate experiment (Dachner & Warren, 2019), we confirmed that visual occlusion reduces heading responses by manipulating the amount of occlusion.

The evidence thus reveals that the neighborhood of interaction results from two decay rates. But what are the underlying processes that produce them? We propose that the gradual decay to nearest (unoccluded) neighbors follows from Euclid’s law of perspective, while the more rapid decay within the crowd is due to the added effect of occlusion. These findings led to a new visual model.

### Visual model

To build a visual model of collective motion from the bottom up, we begin with the visual coupling between a pedestrian and a single neighbor (Bai & Warren, 2019; Dachner & Warren, 2017; Rio et al., 2014).

#### Heading control

Consider a pedestrian who is walking with a neighbor directly ahead of them (eccentricity=0°) (see Figure 4). If the neighbor turns left, this generates a leftward angular velocity in the pedestrian’s field of view (Figure 4a); the pedestrian could cancel the angular velocity by steering left (and vice versa for a right turn). On the other hand, if the neighbor is on the pedestrian’s right (eccentricity=90°) and turns left, this generates an optical expansion in the field of view (Figure 4b); in this case, the pedestrian could cancel the expansion by steering left. Conversely, a right turn by the neighbor generates an optical contraction, which could be canceled by steering right.

**Figure 4.**
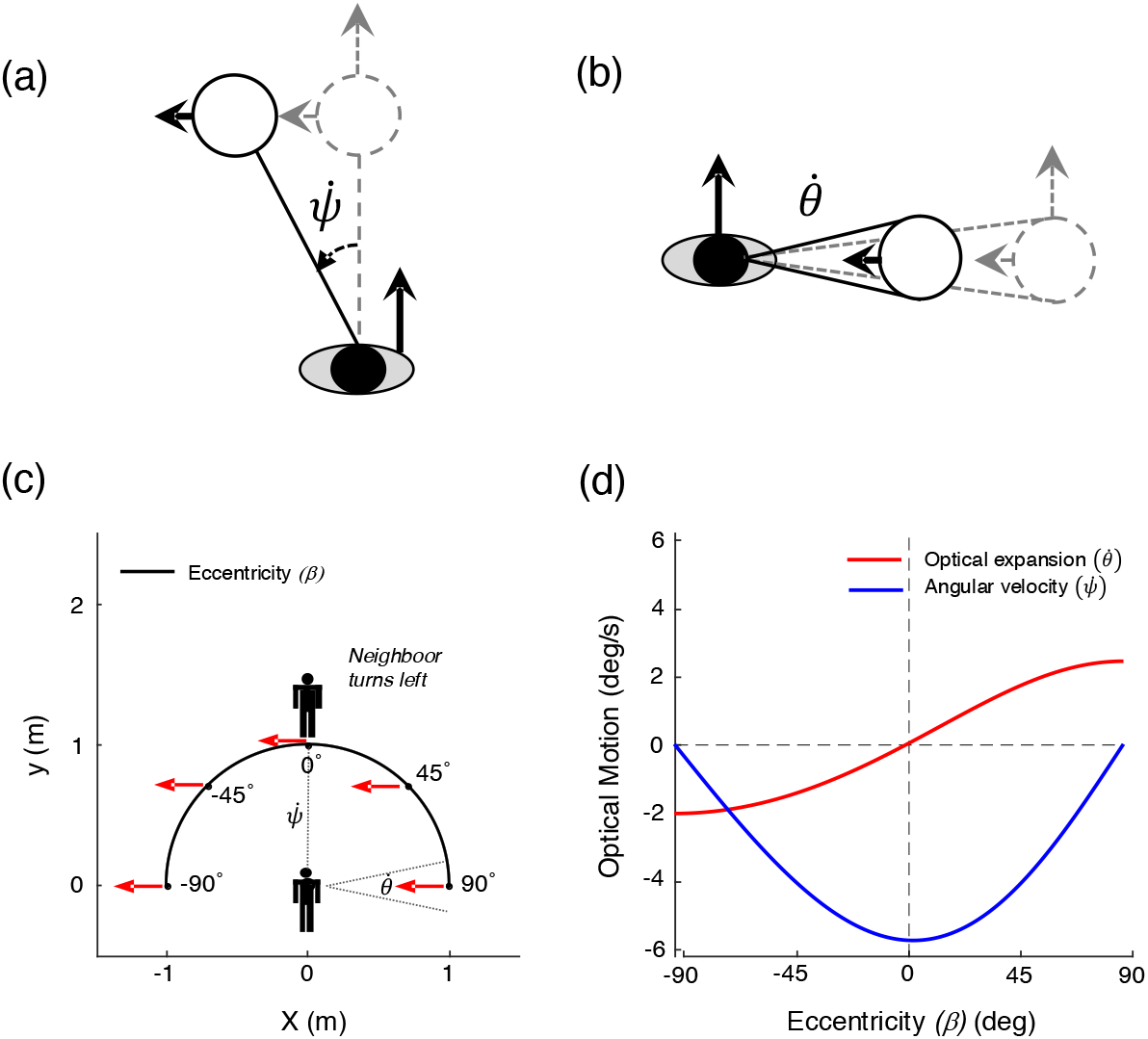
Visual information for the control of heading (walking direction). (a) If a neighbor (open circle) directly ahead of the pedestrian (filled oval) turns left, this creates a leftward angular velocity 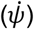 in the pedestrian’s field of view, which the pedestrian can cancel by turning left. (b) If a neighbor on the right side of the pedestrian turns left, this creates an optical expansion 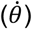 in the field of view, which the pedestrian can also cancel by turning left. (c) The optical motions thus depend on the neighbor’s eccentricity (*β*, black curve), illustrated by a top-down view of a neighbor at a distance of 1m, turning 90° to the left. (d) The resulting optical motion (deg/s) as a function of eccentricity (*β*): Angular velocity (blue curve) is a cosine function of eccentricity, with zero-crossings at *β* = ±90° and a minimum at *β* = 0° (leftward motion). In contrast, expansion rate (red curve) is a sine function of eccentricity, with a zero-crossing at *β* = 0°, a maximum at *β* = 90° (expansion), and a minimum at *β* = −90° (contraction). Note that if the neighbor turns right, these curves are flipped about the x-axis.

These two optical variables thus trade off as a function of the neighbor’s eccentricity (*β*) (Figure 4c,d): If the neighbor turns left, their angular velocity 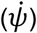 is a cosine function of eccentricity with a minimum (leftward velocity) at *β* = 0°. In contrast, their rate of expansion 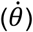 is a sine function with a maximum at *β* = 90° (the minimum at *β* = −90° corresponds to contraction). For a right turn, the sine and cosine functions simply flip about the x-axis. Critically, both optical velocities 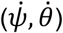 decrease with neighbor distance in accordance with Euclid’s law of perspective, which states that the visual angle subtended by an object, or a motion, diminishes with distance from the observer (specifically, as *tan*^-1^(1/*d*)).

The visual coupling for heading (*ϕ*) can thus be formalized as a second-order control law,

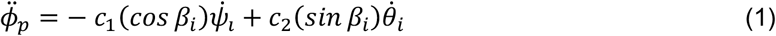

in which pedestrian *p* steers (angular acceleration 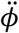) so as to cancel the combined angular velocity 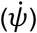 and expansion rate 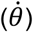 of neighbor *i*. The dependence of these variables on eccentricity (*β*) acts as a filter so the pedestrian is only influenced by components of angular velocity and expansion/contraction that specify a turning neighbor. The free parameters (*c*_1_ = 14.38, *c*_2_ = 59.71) were fit to our previous data on pedestrian following in dyads (Dachner & Warren, 2017) and held constant.

#### Speed control

The control of walking speed is complementary to the control of heading (see Figure 5). If a neighbor directly ahead (*β* = 0°) slows down, this generates an optical expansion in the pedestrian’s field of view (Figure 5a); and if the neighbor speeds up, it generates an optical contraction. But if a neighbor to the pedestrian’s right slows down (*β* = 90°), this generates a negative (leftward) angular velocity (Figure 3b), and vice versa. The two optical variables again trade off as a function of eccentricity, but with the opposite the sine and cosine functions (Figure 5c,d). Once again, by Euclid’s law, the optical velocities decrease with neighbor distance.

**Figure 5.**
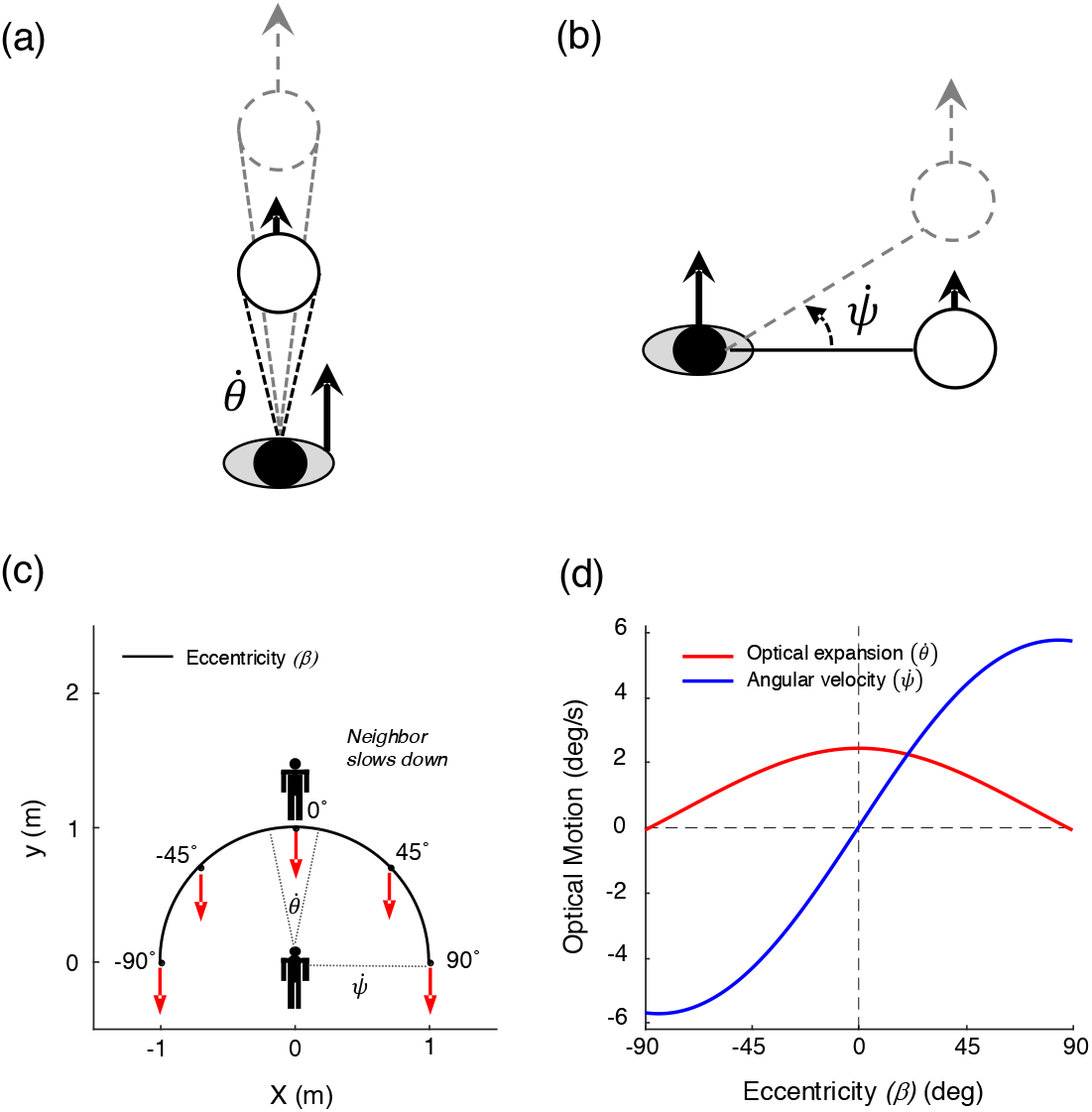
Visual information for the control of walking speed. (a) If a neighbor directly ahead (open circle) slows down, this creates an optical expansion 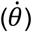 in the pedestrian’s field of view, which the pedestrian (filled oval) can cancel by decelerating. (b) If a neighbor to the pedestrian’s right slows down, this creates an angular velocity 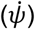 in the field of view, which can also be cancelled by decelerating. (c) The optical motions depend on the neighbor’s eccentricity (*β*, black curve), illustrated by a neighbor at a distance of 1m slowing down by −0.1 m/s. (d) In this case, the resulting angular velocity (blue curve) is a sine function of eccentricity, with a zero-crossing at *β* = 0°, a minimum at *β* = −90° (leftward motion), and a maximum at *β* = 90° (rightward motion), whereas expansion rate (red curve) is a cosine function of eccentricity, with zero-crossings at *β* = ±90° and a maximum at *β* = 0° (expansion). If the neighbor speeds up, these curves flip about the x-axis.

The visual coupling for radial speed 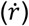 is thus based on the same two optical variables as Equation 1, but the sine and cosine functions are reversed:

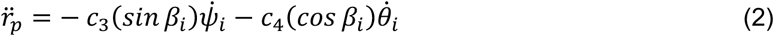

Pedestrian *p* thus linearly accelerates or decelerates 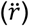 so as to cancel the combined angular velocity 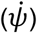 and expansion rate 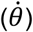 of the neighbor *i*, but the pedestrian is now influenced only by combinations that specify a change in neighbor speed. The free parameters (*c*_3_ = 0.18, *c*_4_ = 0.72) were fit to our data on following in pedestrian dyads (Dachner & Warren, 2017) and held fixed. To account for variation in neighbor size, the relative rate of expansion 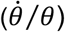 can be substituted for the expansion rate 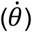, thereby normalizing it by the neighbor’s visual angle (*θ*) (Bai & Warren, 2019; Wagner, 1982).

#### Collective motion

To formulate a visual model of collective motion, we substitute the visual control laws for local interactions (Equations 1 and 2) into a neighborhood function, which averages the influences of multiple neighbors. This yields,

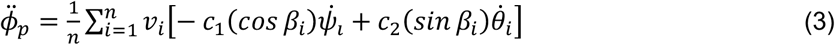

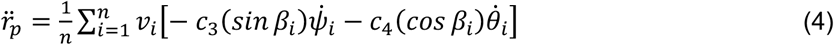

in which pedestrian *p*’s heading and speed are controlled by canceling the mean angular velocity 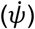 and rate of expansion 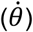 of all visible neighbors (*n*), where the contributions of these two variables trade off as opposite functions of eccentricity (*β*). The 180° field of view is centered on the heading direction, as people tend to face in the direction they’re walking (Grasso, Prevost, Ivanenko, & Berthoz, 1998).

The effect of partial occlusion is incorporated by weighting each neighbor in proportion to their visibility (Dachner & Warren, 2019), ranging from *v_i_* = 0 (fully occluded) to *v_i_* = 1 (fully visible). Visibility is set to 0 if its value falls below a threshold (*v_t_* = 0.15), so *n* is the number of visible neighbors above threshold.

Importantly, due to the geometry of occlusion, the occluded region behind a near neighbor grows with distance, so the visibility of far neighbors tends to decrease with their separation in depth from near neighbors. Consequently, the range of the neighborhood varies with the crowd’s *opacity*, the distance at which farther neighbors are completely occluded (Figure 1b). Note that visual occlusion invalidates the usual assumption of superposition, because the response to far neighbors is not in fact independent of the positions of near neighbors.

Basic properties of previous physical models fall out naturally from the visual model. First, canceling optical expansion yields collision avoidance without an explicit ‘repulsion’ force. Second, canceling optical contraction serves to maintain group cohesion without an explicit ‘attraction’ force. Third, canceling the combined angular velocity and expansion/contraction generates collective motion without an explicit ‘alignment’ rule. Critically, the fact that these variables obey the laws of optics explains the form of the neighborhood. Specifically, Euclid’s law – the diminution of optical velocity with distance – accounts for the gradual decay of influence to the nearest (unoccluded) neighbors in a crowd, and the added effect of occlusion accounts for the more rapid decay within the crowd. The laws of optics thus explain the distance-dependent neighborhood of interaction, without an explicit decay function.

### Simulation results

We tested the visual model by predicting the human trajectories in our virtual crowd experiments and comparing the results to our previous behavioral model (Equations S1-S3). In addition, to assess the contribution of visual occlusion to human responses, we compared the visual model to a *motion model* that was identical, except that the effect of occlusion was removed so responses were based on the optical motion variables alone (see SI for details). We find that the visual model outperforms the behavioral model and generalizes to real crowd data. It also outperforms the motion model, confirming the essential role of visual occlusion.

To simulate each experimental trial, the models were initialized with the participant’s position, heading, and speed 2s before the perturbation. For the behavioral model, the input on each time step was the position, heading, and speed of all virtual neighbors in a 90° field of view. (If an experiment did not manipulate speed, we used the participant’s recorded walking speed from each trial as input to the behavioral model.) For the visual model, the input was the angular velocity, expansion rate, eccentricity, and visibility of each neighbor, calculated from their positions relative to the model on each time step. The output of both models was the position, heading (and speed) of the simulated agent on the next time step, represented as time series for each trial. As a measure of model performance, we then computed the root mean squared error (RMSE) between each participant’s mean time series in each condition and the corresponding mean time series for the model. To compare the two models, a Bayes Factor (BF_VB_) was used to estimate the relative strength of evidence.

#### Simulating two decay processes

First we simulated our double-decay experiment with the visual model. To compare it with the behavioral model, we needed to add a second exponential term to that model’s decay function (Equation S4), which was estimated from the experimental data. The mean final heading for the two models is plotted in Figure 3b, together with the human results. Although both models are close to the 95% confidence intervals for the human data (shaded regions), the visual model (dotted curves) lies entirely within them.

Over the whole time series, the mean heading error for the visual model (RMSE=2.47°) was significantly smaller than that for the behavioral model (RMSE=3.45°) (*t* (9)=14.48, *p*<.001, Cohen’s *d*=1.460), with decisive evidence in favor of the visual model (BF_VB_>>100). The mean position error for the visual model (RMSE=0.241m) was also smaller than that for the behavioral model (RMSE=0.309m) (*t* (9)=8.46, *p*<.001, Cohen’s *d*=0.294), also decisive evidence for the visual model (BF_VB_>>100). A similar comparison between the visual model and the motion model yielded decisive evidence for a contribution of occlusion (BF_VM_>100) (see Figure S1).

In sum, the visual model predicted the distance-dependent neighborhood more closely than the behavioral model, accounting for the two decay processes without an explicit decay function. The visual model thus grounds the neighborhood of interaction in the laws of optics.

#### Re-simulating Rio, Dachner & Warren (2018)

As a further test of the models, we re-simulated the main experiment of Rio, Dachner, and Warren (2018), which perturbed heading or speed and manipulated the number and distance of perturbed neighbors (Figure 6a). Briefly, a participant (n=10) walked with a virtual crowd of 12 neighbors, 5 in the near row (1.5m distance) and 7 in the far row (3.5m distance). On each trial a subset of neighbors (0, 3, 6, 9, or 12, predominantly in the near or far row) turned 10° left or right, or changed speed by ±0.3 m/s from the base speed (1.0 m/s). The participants’ mean final heading and mean final speed appear in Figure 6b,c (solid curves). Responses were larger when near neighbors were perturbed (blue) than when far neighbors were perturbed (red), indicating a decay of influence with distance.

**Figure 6.**
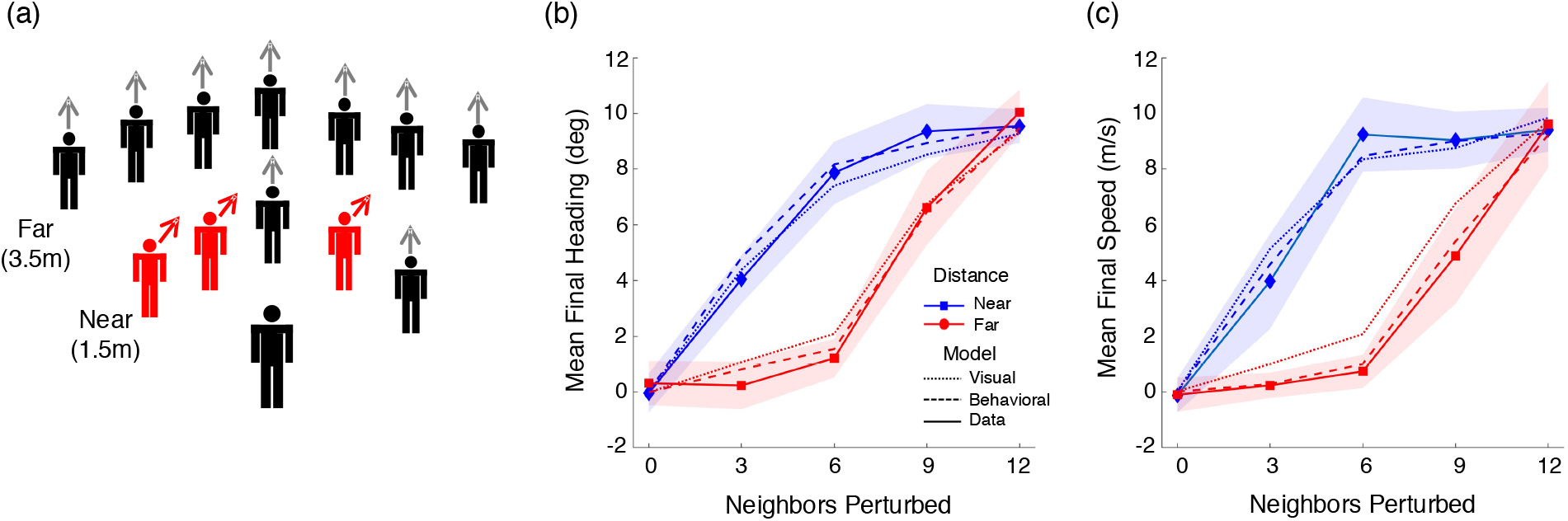
Re-simulation of Rio, Dachner & Warren’s (2018) experiment. (a) Schematic of virtual crowd (n=12), illustrating a subset of neighbors whose heading is perturbed to the right. The number of perturbed neighbors in the near row (1.5m) or the far row (3.5m) was manipulated. Speed was also perturbed in a separate block of trials. (b) Results for heading perturbations: Mean final heading as a function of the number of perturbed neighbors; curves represent the primary perturbed row. (c) Results for speed perturbation: Mean final speed as a function of same. Solid curves correspond to human data, dotted curves to the visual model, dashed curves to the behavioral model with the single-decay function. Shaded regions represent 95% confidence intervals for the human data. [Panels b and c are modified from (8), with permission.]

Simulations of the visual model (dotted curves) and the behavioral model with the original single-decay function (dashed curves) are both close to the human data in Figure 6b,c, falling within the 95% confidence intervals in nearly all conditions. The mean heading error was significantly smaller for the visual model than the behavioral model (RMSE_V_=1.97°, RMSE_B_=2.08°), (*t* (9)=6.94, *p*<.001, Cohen’s *d*=0.871, BF_VB_>100), although there were no differences for the mean speed error (RMSE_V_=0.0627 m/s, RMSE_B_=0.0640), (*t* (9)=1.15, *p*=0.281, Cohen’s *d*=0.208, BF_01_=1.91), or the mean position error (RMSE_V_=0.193m, RMSE_B_=0.199m), (*t* (9)=1.112, *p*=0.295, Cohen’s *d*=0.082, BF_01_=1.96). Both models capture the human data quite well, although the evidence favors the visual model.

The comparatively good performance of the behavioral model in this experiment stems from the fact that the single-decay function was originally fit to human swarms with a density similar to that of the virtual crowd, with nearest neighbors 1-3m distant. However, this empirical decay term does not generalize to larger distances in our double-decay experiment, whereas the visual model does, thanks to Euclid’s law.

Do the observed RMSE values for the visual model indicate good performance? To benchmark the lower bound on model error, we estimated the inherent noise in the data due to gait oscillations and tracker error by computing the mean RMSE between a heading of 0° and the mean time series for each participant in no-perturbation *control trials*. This yielded RMSE_L_=1.53°, revealing that the visual model (RMSE_V_=1.97°) is only 0.44° from the best possible performance. Conversely, to estimate the upper bound on error – a failure to respond to the input – we computed the mean RMSE between a heading of 0° and the mean time series for each participant on *perturbation trials*. This yielded RMSE_U_=4.96°, indicating that the visual model (RMSE_V_=1.97°) is much better than doing nothing. For speed, the visual model (RMSE_V_=0.0627 m/s) is only 0.036 m/s from the best possible performance (RMSE_L_=0.027 m/s) and again is much better than doing nothing (RMSE_U_=0.113 m/s).

In sum, the visual model accounts for Rio, et al.’s (2018) experiment as well or better than the behavioral model. Although the latter *describes* the relation between neighbor and participant velocities, the visual model *explains* that relation by appealing to the laws of optics: neighbors in the far row exert less influence because they have lower optical velocities due to Euclid’s law and are partially occluded by near neighbors. We confirmed the role of occlusion by re-simulating the experiment with the motion model (without occlusion), yielding decisive evidence for the visual model (BF_VM_>100) (Figure S2).

#### Human swarm

To test whether our findings for virtual crowds apply to real crowds, we recorded the walking trajectories of pedestrians in ‘human swarms’, intended to mimic collective motion in a public space. A group of participants was instructed to walk about a large hall, veering left and right, while staying together as a group. We then attempted to predict the trajectory of an individual participant from the movements of their neighbors, using the visual and behavioral models.

We recorded three different groups (n=10, 16, 20) walking together in a large tracking area (14m x 20m) for 2 min trials. Density was manipulated by varying the size of a starting box marked on the floor, and each group performed two trials at each density (measured as low=1.72 and high=2.10 participants/m^2^), for a total of 12 trials. Head-mounted markers were tracked using 16 motion-capture cameras (60 Hz) and time series of position, heading and speed were computed as before. We then identified 10s segments of data in which ≥75% of the participants were continuously tracked, yielding 30 segments for analysis (17 high and 13 low density). In each segment, we simulated a focal participant at the back of the group and treated the tracked neighbors as input on each time step. We used the original single-decay function in the behavioral model (Equation S3), which had been fit to a sample of the swarm data.

Two segments of simulated swarm data appear in Figure 7. The heading time series (b) for the focal participant (red) is more closely captured by the visual model (blue) than the behavioral model (green) in both segments, whereas the speed time series (c) is better approximated by the behavioral model in Segment 1 and the visual model in Segment 10 (panel c). Over all 30 segments, the mean heading error was significantly lower for the visual model (RMSE_V_=15.0°) than the behavioral model (RMSE_B_=22.9°) (*t* (29)=4.48, *p*<0.001, Cohen’s *d*=0.806), decisive evidence for the visual model (BF_VB_>100), and so was the mean position error (RMSE_V_=0.60m, RMSE_B_=0.80m) (*t* (29)=2.21, p<0.05, Cohen’s *d*=0.338), anecdotal evidence (BF_VB_=1.60). On the other hand, the mean speed error was significantly lower for the behavioral model (RMSE_V_=0.224 m/s, RMSE_B_=0.146 m/s) (*t* (29)=6.83 *p*<0.001, Cohen’s *d*=1.198), decisive evidence (BF_BV_>>100), although variation in speed accounted for less of the total variation in position. Finally, the visual model performed better than the motion model (without occlusion) (BF_VM_>40 or better), confirming the importance of occlusion (Figure S3).

**Figure 7.**
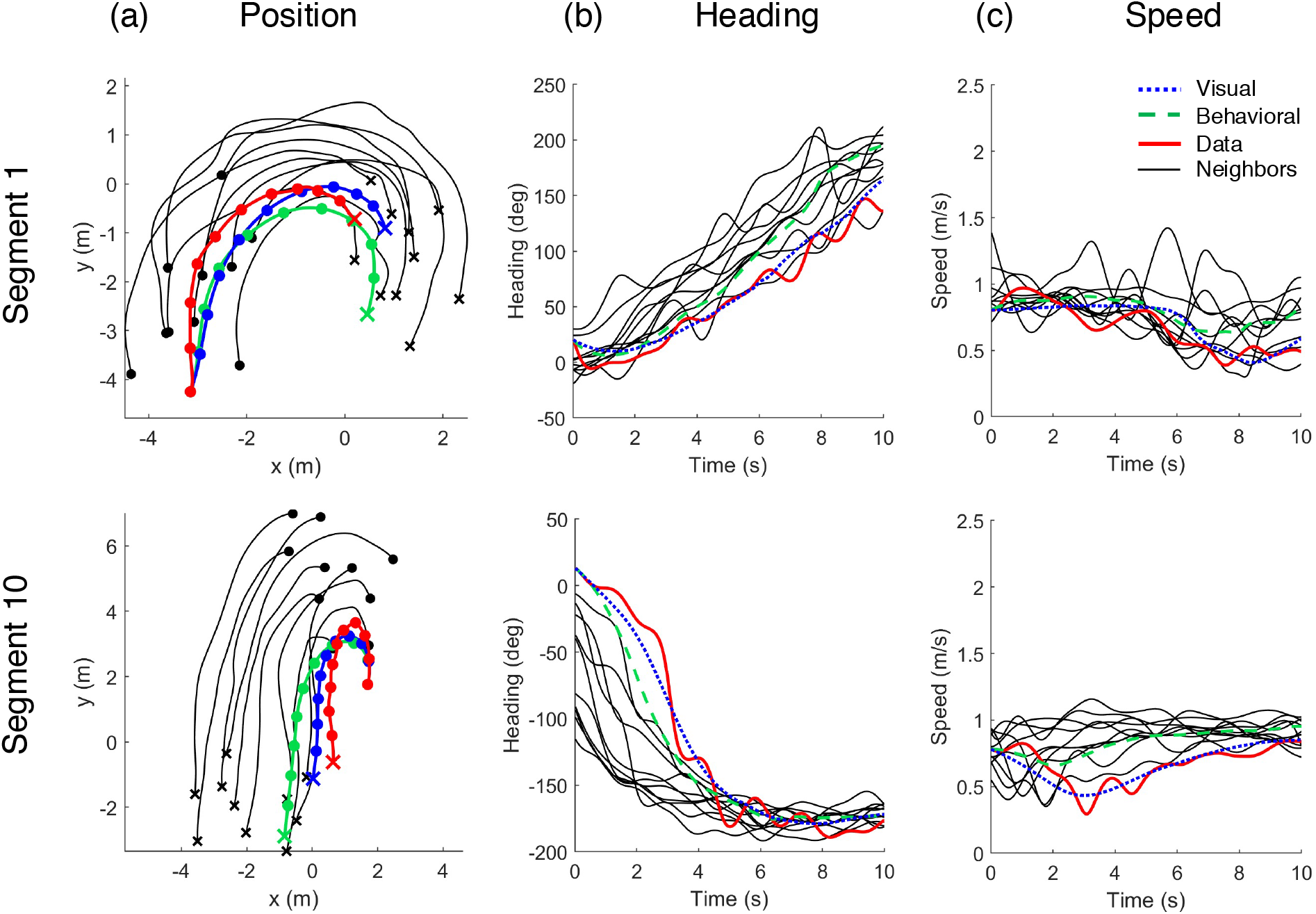
Sample data from the human swarm. Each row represents a 10s segment of data, with the focal participant (red) and simulations of the visual model (blue) and the behavioral model (green). (a) Traces of position over time (Segment 1: RMSE_V_ = 0.379m, RMSE_B_=0.818; Segment 10: RMSE_V_=0.275m, RMSE_B_=1.389m). Thin gray curves = tracked neighbors that were input to the model; o = starting positions, x = final positions, dots at 1s intervals. (b) Time series of heading (Segment 1: RMSE_V_=10.67°, RMSE_B_=32.88°; Segment 10: RMSE_V_=11.81°, RMSE_B_=23.62°). (c) Time series of speed (Segment 1: RMSE_V_=0.187 m/s, RMSE_B_=0.162 m/s; Segment 10: RMSE_V_=0.157 m/s, RMSE_B_=0.178 m/s). Note that errors in the swarm are higher than those for virtual crowds because they represent simulations of single trials rather than subject mean time series, and thus reflect gait oscillations and tracking errors.

Overall, the visual model accounts for individual trajectories in real crowd data better than the behavioral model, even though the latter’s decay term was fit to a subset of the same data. We attribute this advantage primarily to the effect of visual occlusion. Whereas the behavioral model approximates the decay with distance using a fixed exponential decay function, the visual model incorporates the dynamic occlusion that is visible on each trial, and is thus sensitive to the variation in occlusion over time.

## Discussion

Nearly all models of collective motion in humans and animals attribute local interactions to hypothesized rules or metaphorical forces, described from an overhead view. In this article we developed a bottom-up visual model of human ‘flocking’, grounded in the optical information known to govern pedestrian interactions from an embedded viewpoint. In contrast to previous descriptive models, the visual model explains basic properties of interaction as a natural consequence of the laws of optics.

First, *rules and forces of interaction* are reduced to optical variables that control an individual’s heading and speed. Instead of explicit ‘repulsion’ and ‘attraction’ forces, canceling optical expansion results in collision avoidance, while canceling optical contraction maintains group cohesion. Instead of an explicit ‘alignment’ rule, collective motion emerges from canceling the combined expansion/contraction and angular velocity of neighbors. This visual coupling yields a robust response regardless of the individual’s position within the group.

Second, the *neighborhood of interaction* is explained by the laws of optics, without an explicit distance term. The gradual decay to the nearest (unoccluded) neighbors in a crowd follows from Euclid’s law, the diminution of optical velocity with distance. The more rapid decay within a crowd follows from the added effect of visual occlusion, which grows with the separation in depth between near and far neighbors. Consequently, the neighborhood range and number of neighbors *n* do not depend on a fixed metric or topological distance (Ballerini et al., 2008; Strandburg-Peshkin et al., 2013), but vary with crowd opacity.

Interestingly, the visual model predicts that the effective neighborhood depends on crowd density, which we have confirmed in other experiments (Wirth, Dachner, Rio, & Warren, 2021; Wirth & Warren, 2018). In dense human crowds (1-2m apart), complete opacity can occur by 5m, whereas lower densities yield larger ranges of interaction. In the limit, starlings appear to adjust flock density to maintain a ‘marginal opacity’ at which individual birds can see through the entire flock(Pearce, Miller, Rowlands, & Turner, 2014). The range of interaction might also be limited by a detection threshold for optical motion. However, adding a motion threshold to the visual model did not improve the fit to the data, perhaps because it was superseded by complete opacity.

Virtually all physical models assume the principle of superposition for combining binary interactions between individuals. However, the superposition principle is invalidated by the facts of visual occlusion. Because the influence of a far neighbor depends on the position of a near neighbor, an agent’s response is not a linear combination of independent responses to each neighbor. While this may be computationally inconvenient, visual occlusion has large effects on local interactions and any model should take them into account.

Note that Euclid’s law predicts an asymmetry in a pedestrian’s response. Assume a neighbor at a given initial distance ahead: if they slow down, their distance decreases; whereas if they speed up by the same amount, their distance increases. Consequently, the corresponding rate of expansion is greater than the rate of contraction, respectively. This effect explains an asymmetric speed response we previously observed in pedestrian following (Bai & Warren, 2019; Rio et al., 2014).

We found that the visual model generally outperforms the behavioral model, although they were quite similar in our re-simulation of Rio, Dachner & Warren’s (2018) data. This is attributable to the fact that the behavioral model approximates the effect of distance with an exponential decay term that was fit to human swarms that had a density similar to the virtual crowd display. However, this decay term did not generalize to larger crowd distances in our double-decay experiment, whereas the visual model did, with fixed parameters. The visual model thus explains the form of the neighborhood and generalizes to new conditions without re-parameterization.

We noted a limitation of the current visual model when simulating the swarm data. In five additional segments, the front of the crowd executed a 180° hairpin turn and walked back toward the focal participant, generating a rapid expansion in the field of view. Whereas human participants kept walking forward, the visual model responded by slowing down and backing up to cancel the optical expansion. Similar but less extreme responses to U-turns may explain the higher speed RMSE for the visual model reported above. Clearly, the model needs to distinguish neighbors in a group from obstacles to be avoided, which may be as straightforward as distinguishing the face and back of other pedestrians.

Our findings suggest that characteristic patterns of collective motion in different species might result from reliance on different sensory variables. Humans cancel optical velocities, which will yield collective motion despite variation in the distance, density, and size of neighbors. In contrast, holding the visual angles of neighbors at a particular value would yield a school with a preferred spatial scale, whereas maintaining neighbors in particular visual directions would yield a flock with a preferred spatial structure (Ballerini et al., 2008).

In sum, we conclude that the local interactions underlying collective motion have a lawful basis in the visual coupling between neighbors. We are pursuing the consequences of this finding for other aspects of collective crowd motion, including information transfer, leadership networks (Lombardi, Warren, & di Bernardo, 2020), and the role of visual attention (Lemasson, Anderson, & Goodwin, 2009).

## Methods

### Experimental methods

#### Human subjects

Participants were undergraduate and graduate students at Brown University, with normal or corrected-to-normal vision and none reported having a motor impairment. Different individuals participated in each experiment. A power analysis for repeated-measures ANOVA determined that to achieve a power of 1 – *β* = 0.85 with an error probability of *α* =.05 and an effect size of 0.5 (*η*^2^ = 0.2) required a sample size of 8 per experiment. The research protocol was approved by Brown University’s Institutional Review Board, in accordance with the principles expressed in the Declaration of Helsinki. Informed consent was obtained from all participants, who were paid for their participation.

#### Equipment

VR experiments were conducted in the Virtual Environment Navigation Lab (VENLab) at Brown University. Participants walked freely in a 12m x 14m tracking area while viewing a virtual environment in a wireless stereoscopic head-mounted display (HMD) (Rift DK1, Oculus, Irvine CA; 90°H x 65°V field of view, 640 x 800 pixels per eye, 60 Hz refresh rate). Head position and orientation were recorded with a hybrid inertial/ultrasonic tracking system (IS-900, Intersense, Billerica MA; 60 Hz sampling rate), and used to update the display with a total latency of 50-67 ms.

#### VR displays

The virtual environment was generated in Vizard software (WorldViz, Santa Barbara, CA). A green start pole and a gray orienting pole (radius 0.2m, height 3m) appeared 12.73 m apart on a ground plane with a grayscale granite texture and a blue sky. Virtual humans (WorldViz Complete Characters) were animated 3D models (M=8073 ±780 polygons, each polygon with 2048 x 2048 pixels). They were initially placed on arcs with the participant’s start pole at the center, at equally spaced eccentricities (±6°, ±19°, ±32°, or ±45°) from the participant’s initial walking direction toward the orientation pole. These initial positions were then randomly jittered in polar coordinates, within an angular range of 5° and a radial range of 0.2m. A different crowd configuration was generated for each trial. The virtual humans were animated with a walking gait with randomly varied phase (updated at 60 Hz), and the crowd contained equal numbers of men and women and diverse races and ethnicities. During a trial, all virtual humans accelerated from a standstill (0 m/s) to a speed of 1.0 m/s on a straight path over a period of 3s, following a sigmoidal function (cumulative normal, *μ* = 0, *σ* = 0.5 s) fit to previous human data. They continued to travel with the same heading and speed for an additional 2s. Then the heading of some or all virtual humans was perturbed by changing their walking direction over a period of 0.5s following a similar sigmoidal function; the display continued for another 7s.

#### Procedure

Participants were instructed to “walk with the group of virtual humans” and to “treat them as if they were real people.” On each trial, the participant walked to the green start pole and faced the gray orienting pole. After 2s, the poles disappeared and the virtual crowd appeared; 1s later, a verbal command (“Begin”) was played over the HMD’s headphones and the crowd began walking. The display continued until the participant had walked for 12s, and a verbal command (‘End’) signaled the end of the trial. The start pole for the next trial then appeared nearby. Test trials were preceded by two practice trials to familiarize the participant with walking in the virtual environment.

#### Data processing

Data were processed in Matlab (Mathworks, Natick, MA). For each trial, the time series of head position in the horizontal (X–Y) plane were filtered using a forward and backward fourth-order low-pass Butterworth filter to reduce tracker error and oscillations due to the step cycle. Time series of heading direction and walking speed were then computed from the filtered position data. A 0.6 Hz cut-off was used before computing heading to reduce lateral oscillations on each stride, while a 1.0 Hz cutoff was used before computing speed to reduce anterior–posterior oscillations on each step. The dependent measure was the *final heading*, the average heading direction during the last two seconds of each trial. Final heading responses that were more than 3 standard deviations away from that individual subject’s mean were removed from the data set, then the mean final heading was computed for each participant in each condition.

#### Statistical analysis

Statistics were computed in Excel (Microsoft) and JASP (https://jasp-stats.org/). A preliminary 3-way Repeated-Measures ANOVA on final heading (Factor A x Factor B x left/right/control) was performed to confirm that the heading manipulation was successful in both directions. Left/right trials were then collapsed for further analysis by multiplying the final heading on left trials by −1. The collapsed data were analyzed using 2-Way Repeated Measures ANOVA, followed by simple-effects tests or Bonferroni-corrected post-hoc *t*-tests (two-tailed). Effect sizes for ANOVAs were estimated using general eta-squared 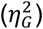; effect sizes for paired *t*-tests were estimated using Cohen’s *d*, conservatively computed as the difference of the two means divided by the pooled variance of the two SDs (Dunlap, Cortina, Vaslow, & Burke, 1996).

### First experiment: Decay to nearest neighbors

#### Participants

Twelve participants (7 female, 5 male).

#### Displays

The crowd consisted of 2, 4, or 8 virtual humans, initially placed on one arc (radius = 1.8, 3.0, 4.0, 6.0 or 8.0 m), at randomly assigned eccentricities (±6°, ±19°, ±32°, or ±45°). The heading direction of all virtual humans was perturbed to the right (+10°) on one-third of the trials, and to the left (−-10°) on one-third of the trials; on the remaining control trials, they did not turn but continued walking on a straight path (0°).

#### Design

5 crowd distances (1.8, 3.0, 4.0, 6.0, 8.0 m) x 3 crowd sizes (2, 4, 8 neighbors) yielded 15 conditions. Each participant received 4 heading perturbation trials (±10°) and 2 control trials (0°) per condition, for a total of 90 trials. They were presented in a randomized order in a 1-hour session.

### Second experiment: Double-decay hypothesis

#### Participants

Ten participants (6 female, 4 male) successfully completed Experiment 2; two additional participants discontinued the experiment due to discomfort.

#### Displays

The crowd consisted of 12 virtual humans, with four of them initially positioned on three concentric arcs (near, middle, far rows), with the start pole at the center. The arcs were separated by 2m, and radius of the near arc was varied (crowd distance = 2, 4, or 6 m). The heading direction of all four neighbors in one row was perturbed to the right (+10°) on one-third of the trials, and to the left (−10°) on one-third of the trials; on the remaining one-third of trials (control trials), there was no perturbation (0°) and all neighbors walked on a straight path.

#### Design

3 crowd distances (2, 4, or 6 m) x 3 perturbed rows (near, middle, far) yielded 9 conditions. Each participant received 6 heading perturbation trials (±10°) and 3 control trials (0°) per condition, for a total of 81 trials, presented in a randomized order in a 1-hour session.

Note that heading responses were expected to be smaller than in the first experiment because only one-third of the crowd was perturbed on each trial.

### Human swarm

#### Participants

Three different groups (N=10 (5F, 5M), N=16 (6F,10M), N=20 (10F,10M)) participated in separate sessions as part of a larger study.

#### Equipment

Head position was recorded in a large hall with a Qualisys Oqus 16-camera infrared motion capture system (Qualisys, Buffalo Grove, IL) at 60Hz. The tracking area (14m x 20m) and starting boxes of various sizes were marked on the floor with colored tape. Each participant wore a bicycle helmet with a unique constellation of five reflective markers on 30–40 cm stalks.

#### Procedure

Participants were instructed to walk about the tracking area at a normal speed for periods of 2 min, veering randomly left and right, while staying together as a group. Each group received two trials in each of two density conditions (low, high), determined by the size of the starting box. Participants began each trial in shuffled positions in a specified box; at a verbal ‘go’ signal, they began walking for 2 min, until a ‘stop’ signal. This yielded mean densities of 1.72 participants/m^2^ in the low density condition and 2.10 participants/m^2^ in the high condition, as measured within the bounding polygon of the crowd over all data (using the Matlab *boundary* and *polyarea* functions).

#### Design

3 groups (N=10, 16, 20) x 2 densities (low, high) with 2 trials in each condition, yielding a total of 12 trials with 24 min of raw data.

#### Data processing

The 3D positions of the markers on each helmet were reconstructed and tracked using QTM software (Qualisys), and the centroid for each helmet was computed using a custom algorithm. Due to infrared reflections in the hall, there were many tracking errors: 100% of the helmets were recovered in only 45% of all frames. The time series of head position in the horizontal (x,y) plane was filtered in Matlab as before, and the heading direction and speed of each helmet was computed on each time step in which it was successfully tracked. We identified 35 10s segments of data in which ≥75% of the participants were continuously tracked. Five of these included hair-pin turns in which the front of the crowd walked back toward the focal participant, creating large optical expansions that caused the model to stop or back up; these were treated separately. This yielded 30 segments (17 high density and 13 low density) in which the positions of most neighbors were known. Rio, Dachner & Warren (2018) previously simulated the same segments with the behavioral model, but subsequent improvements in data processing yielded more tracked neighbors, so the present results are more accurate.

### Simulation methods

#### Simulations of virtual crowd experiments

Individual trials were simulated in Matlab using the Runge-Kutta method (*ode45* function). The participant’s position, heading, and speed 2s before the perturbation were taken as the initial conditions. For the behavioral model (Equations S1-S2), the input on each time step was the position, velocity, and speed of the virtual humans in the participant’s field of view for that trial. The double-decay experiment was simulated using the double-decay function (Equation S4) and Rio, Dachner & Warren’s experiment was simulated using the original single-decay function (Equation S3). Because speed control was not studied in the double-decay experiment, the virtual crowd’s speed was constant, so the recorded time series of the participant’s walking speed during each trial was treated as input to the behavioral model. For the visual model (Equations 3–4), the input was the angular velocity, optical expansion rate, eccentricity, and visibility of each virtual human, which were calculated from their positions relative to the simulated agent on each time step. The output of both models was the position, heading, and speed of the simulated agent on the next time step. This was represented as a time series of heading, a time series of speed, and time series of (x,y) position for every trial.

#### Model comparisons

To compare the simulations with the human data, we calculated the model’s mean time series for each participant in each condition, and then computed the root of the mean squared error (RMSE) between the model mean time series and the corresponding participant mean time series. The overall mean RMSE values for the two models were compared using paired *t*-tests (two-tailed) and scaled JZS Bayes Factors (BF) with a Cauchy prior of 0.707, which indicate the relative strength of evidence for the hypotheses (Rouder, Speckman, Sun, Morey, & Iverson, 2009). Note that the variability in final heading for the models is very small compared to the human data because gait oscillations and tracker error were not simulated; thus, the model means are compared with 95% confidence intervals for the human data in the figures.

#### Model performance benchmarks

The performance of any model is limited by the inherent noise in the human data due to gait oscillations and tracker error. To benchmark the lower bound on error, we estimated the fluctuations around a straight walking direction (heading = 0°) by computing the RMSE_L_ between a value of 0° and each participant’s mean time series of heading on no-perturbation *control* trials. Conversely, to benchmark the upper bound on error, we estimated the error for a model that does not respond to the input (heading = 0°) by computing the RMSE_U_ between a value of 0° and each participant’s mean heading time series on *perturbation* trials. These benchmarks indicate the range of model performance, from the best possible performance given the noise in the data, to the performance of a model that does nothing. (Of course, there exist even worse models that respond inappropriately.)

#### Human swarm simulations

The participant farthest back in the swarm at the beginning of each 10s segment was selected as the focal participant. The focal participant’s position, heading, and speed at the start of the segment were taken as initial conditions for the model simulation. For the behavioral model (Equations S1-S3), the position, heading, and speed of all tracked neighbors were treated as input to the model, using the original single decay function. For the visual model (Equations 3–4), optical variables (angular velocity, expansion rate, eccentricity, and visibility) were computed from the neighbor positions relative to the simulated agent on each time step and used as input. Output was the position, heading, and speed of the simulated agent on the next time step. This was represented as a time series of heading, a time series of speed, and time series of (x,y) position for each segment (e.g. Figure 7). The RMSE between the time series for the simulated agent and the that for the focal participant was computed for each segment, and the mean RMSE for each model was calculated over all 30 segments. Note that the RMSE values were higher than those in the virtual crowd experiments because they represented simulations of a single trial, rather than the mean time series of multiple trials; consequently, they reflected larger fluctuations due to gait oscillations, tracker error, and incomplete tracking of neighbors in the swarm.

## Acknowledgments

This research was supported by the National Institutes of Health, R01EY010923 and R01EY029745 to W.W., and T32 EY018080 to Brown University; the National Science Foundation, BCS-1431406 to W.W.; and Link Foundation Fellowships to G.D. and T.W. Thanks to Adam Kiefer, Stephane Bonneaud, Michael Fitzgerald, and the Sayles Swarm crew for their help during crowd data collection, and to Arturo Cardenas, Eugy Han and the VENLab team for their assistance in processing and analyzing the datafiles.

## Author contributions

G.D. and W.W. conceived the project and designed the research; T.W. and E.R. performed the experiments; T.W. and W.W. analyzed the data; G.D. and W.W. developed the model; G.D. simulated the data and analyzed the results; and G.D. and W.W. wrote the paper.

## Notes

### Competing Interest Statement

The authors have declared no competing interest.

