## Supplementary Information for "The visual coupling between neighbors explains ‘flocking’ in human crowds"

### Model variants

*Behavioral model*

Rio, Dachner, and Warren (2018) described an 'overhead' behavioral model of collective crowd motion based on the alignment of velocity (Figure 1a). The rules of engagement were derived from experiments on following in pairs of pedestrians (Dachner & Warren, 2014; Rio, Rhea, & Warren, 2014), which showed that a pedestrian tends to match the heading direction and speed of a neighbor. The neighborhood of interaction was derived from experiments on a participant walking with a virtual crowd (Rio et al., 2018). Together, this yielded a weighted averaging model of heading control,

$$\ddot{\phi}_p = -\frac{k}{n}\sum_{i=1}^{n} w_i sin\left(\phi_i - \phi_p\right) \quad \text{(S1)}$$

where pedestrian *p*'s angular acceleration (change in heading $\phi$) is proportional to the mean difference between *p*'s current heading and the heading of each neighbor *i*, weighted ($w_i$) by a distance term. The sine function defines an attractor in the neighbors' mean heading direction (a circular variable), and the stiffness or gain *k*=3.15 was fit to data on following in pedestrian dyads (Dachner & Warren, 2014). Here *n* is the number of neighbors within a 5m radius and a 180° field of view, centered on the participant's current heading direction.

Speed is controlled by an analogous equation for *p*'s linear acceleration,

$$\ddot{r}_p = \frac{c}{n}\sum_{i=1}^{n} w_i\left(\dot{r}_i - \dot{r}_p\right) \quad \text{(S2)}$$

which defines an attractor at the neighbors' mean walking speed, weighted by their distance. The constant *c*=3.61 was fit to the pedestrian following data (Dachner & Warren, 2014).

In both equations, the weight $w_i$ of each neighbor decays as an exponential function of distance $d_i$,

$$w_i = \frac{a}{\mathrm{e}^{\omega d}{}_{i+a}} \quad \text{(S3)}$$

where the decay rate $\omega = 1.3$ and scaling constant *a*=9.2 were fit to real crowd data from the human swarm, such that $w_i$ asymptotes near zero at 4-5m(Rio et al., 2018). This yields a soft metric neighborhood of interaction with a gradual decay, which is known to generate robust collective motion in numerical simulations (Cucker & Smale, 2007).

To account for the results of the double-decay experiment, we estimated a double-decay function for the behavioral model by adding a second exponential component to Equation S3:

$$w_i = \left(\frac{a}{e^{\eta(d_i)} + e^{\omega(d_i - d_{NN})} + a}\right) \quad \text{(S4)}$$

where $\eta = 0.4$ is the gradual decay rate to the nearest neighbors,d fit to the experiment data, $\omega = 1.3$ is the previous decay rate within the crowd, and *a*=9.2 is the previous scaling constant.

*Motion model*

To investigate the contribution of visual occlusion to the participant's heading and speed response, we created an *optical motion model* that was identical to the visual model, but removed the effects of partial or full occlusion (i.e. set visibility weight $v_i = 1$). The model thus controls heading and speed by canceling the combined angular velocity ($\dot{\psi}$) and the rate of expansion ($\dot{\theta}$) of all neighbors in the field of view (n), ignoring occlusion. This is equivalent to,

$$\ddot{\phi}_p = \frac{1}{n}\sum_{i=1}^{n}\left[-c_1(\cos\beta_i)\dot{\psi}_i + c_2(\sin\beta_i)\dot{\theta}_i\right] \tag{5}$$

$$\ddot{r}_p = \frac{1}{n}\sum_{i=1}^{n}\left[-c_3(\sin\beta_i)\dot{\psi}_i - c_4(\cos\beta_i)\dot{\theta}_i\right] \tag{6}$$

The parameter values ($c_1 = 14.38, c_2 = 59.71, c_3 = 0.18, c_4 = 0.72$) were the same as for the visual model.

### Supplementary Results

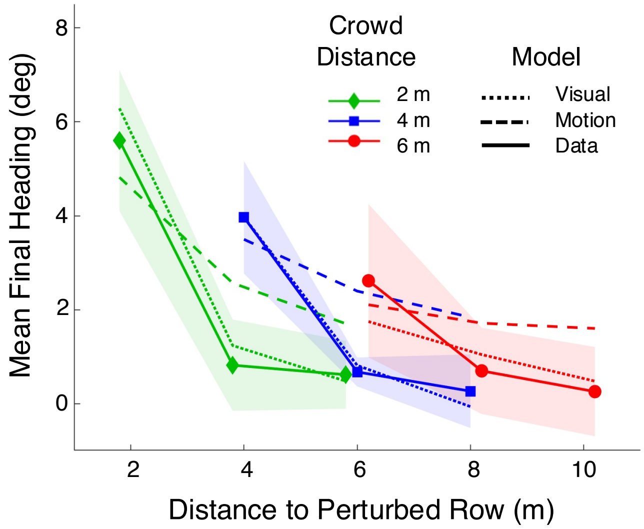


Figure S1. *Double-decay experiment:* To check the contribution of visual occlusion to the heading response, we compared simulations of the visual model (with occlusion, Equation 3) with simulations of the motion model (without occlusion, Equation S5). Mean final heading is plotted as a function of distance to the perturbed row for the human data (solid curves), simulations of the motion model (dashed curves), and simulations of the visual model (dotted curves). Shaded regions are the 95% confidence intervals for the human data.

The visual model is close to the human mean and within the 95% confidence interval in all conditions, whereas the motion model overshoots the human data in most conditions because it is influenced by occluded neighbors. The mean heading error for the visual model ($RMSE_V$=2.47˚) is significantly smaller than that for the motion model ($RMSE_M$=3.28˚), ($t(9)$=41.56, $p$<0.001, Cohen's $d$=1.31), decisive evidence for the visual model ($BF_{VM}$>>100). Similarly, the mean position error for the visual model ($RMSE_V$=0.241m) is significantly smaller than the that for the motion model ($RMSE_M$ = 0.273), ($t(9)$=28.61, $p$<0.001, Cohen's $d$=0.139), indicating very strong evidence for the visual model ($BF_{VM}$>>100). These results indicate that visual occlusion makes an essential contribution to the heading response.

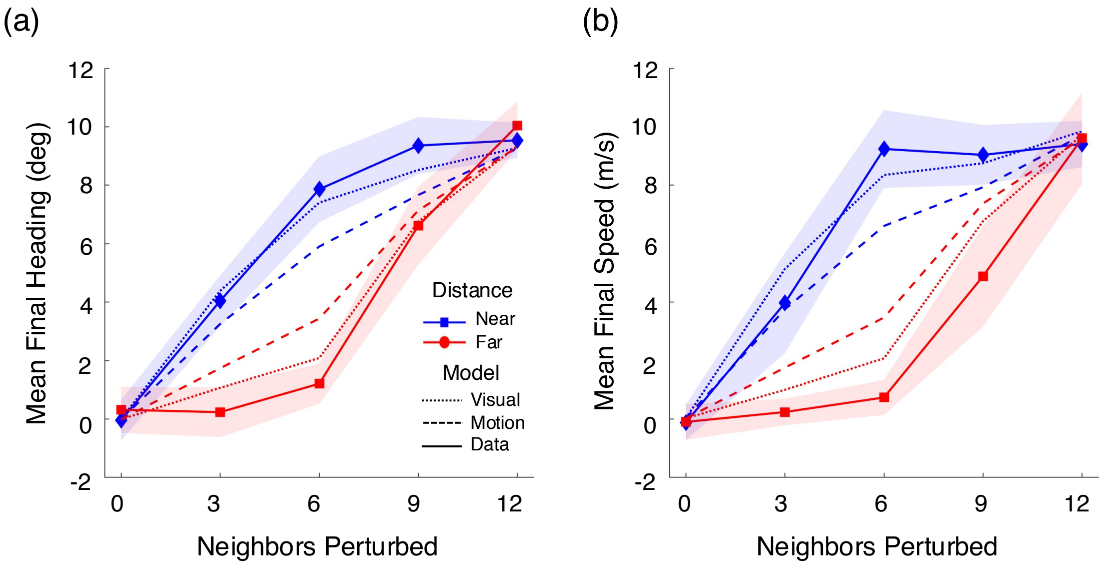


Figure S2. *Rio, Dachner & Warren's experiment:* We also checked the contribution of visual occlusion by re-simulating the data from Rio, Dachner, and Warren *(2018)* with both the visual model (with occlusion, Equations 3-4) and the motion model (without occlusion) (Equations S5-S6). (a) Mean final heading, and (b) mean final speed as a function of the number of neighbors perturbed (out of 12). Solid curves are the human data, dashed curves are simulations of the motion model, and dotted curves are simulations of the visual model. Shaded regions are the 95% confidence intervals for the human data.

When the near row of neighbors is perturbed (blue curves), the motion model undershoots the data, and when the far row is perturbed (red curves) it overshoots the data. However, the visual model more closely captures the data, falling within the human 95% CI in all but one condition. The mean heading error for the visual model ($RMSE_V$=1.97˚) was significantly less than that for the motion model ($RMSE_M$=2.47˚), ($t(9)$=40.06, $p$< 0.001, Cohen's $d$=3.68), with decisive evidence for the visual model ($BF_{VM}$>>100). The mean speed error for the visual model ($RMSE_V$=0.063 m/s) was also significantly smaller than that for the motion model ($RMSE_M$=0.073 m/s), ($t(9)$=13.14, $p$<0.001, Cohen's $d$=1.66), decisive evidence for the visual model ($BF_{VM}$>>100). Similarly, the mean position error for the visual model ($RMSE_V$=0.193m) was significantly smaller than that for the motion model ($RMSE_M$=0.252m), ($t(9)$=78.14, $p$<0.001, Cohen's $d$=0.78) decisive evidence for the visual model ($BF_{VM}$>>100). Thus, when far neighbors were occluded by near neighbors, their influence was reduced and the participant was primarily influenced by near neighbors. These results confirm that occlusion plays a major role in both heading and speed responses.

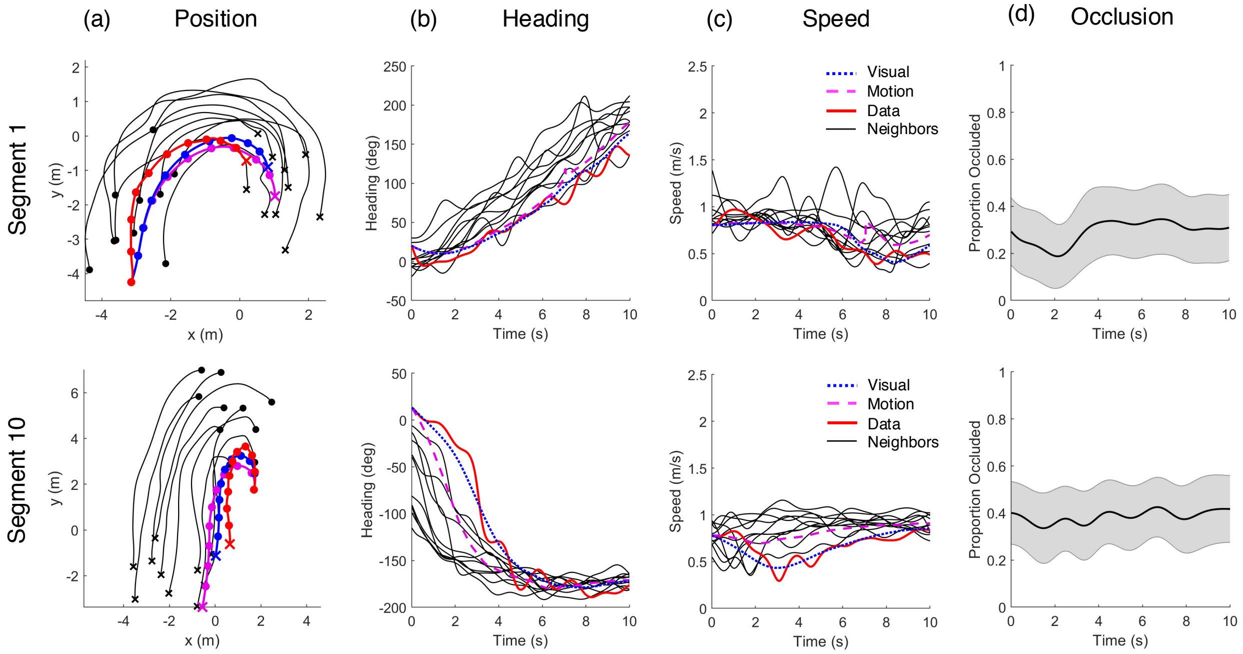


Figure S3. *Human swarm:* We also checked the role of occlusion in the human swarm data. Each row represents a 10s segment of swarm data, with the focal participant (solid red) and simulations of the visual model (dotted blue) and the motion model (dashed magenta). (a) Traces of position over time (Segment 1: $RMSE_V$ = 0.379m, $RMSE_M$=0.772; Segment 10: $RMSE_V$ = 0.275m, $RMSE_M$=1.921m). Thin gray curves = tracked neighbors that were input to the models; o = starting positions, x = final positions, dots at 1s intervals. (b) Time series of heading (Segment 1: $RMSE_V$ = 10.67˚, $RMSE_M$=16.35˚; Segment 10: $RMSE_V$ = 11.81˚, $RMSE_M$=55.85˚. (c) Time series of speed (Segment 1: $RMSE_V$ = 0.187 m/s, $RMSE_M$=0.257 m/s; Segment 10: $RMSE_V$ = 0.157 m/s, $RMSE_M$=0.304 m/s). Note in Segment 1 at 7s the motion model accelerates in response to an occluded neighbor, while the visual model and the participant do not. (d) Time series of the mean proportion of occlusion over all neighbors. Shaded region is 95% confidence interval.

Over all 30 segments analyzed, the mean heading error for the visual model ($RMSE_V$=15.00˚), was significantly smaller than the motion model ($RMSE_M$=32.58˚), ($t(29)=4.41$, $p<0.001$, Cohen's $d=0.96$), as was the mean speed error ($RMSE_V$=0.224 m/s, $RMSE_M$=0.265 m/s), ($t(29)=4.69$, $p<0.001$, Cohen's $d=0.64$), both providing decisive evidence for the visual model ($BF_{VM}>100$). The mean position error was also significantly smaller for the visual model ($RMSE_V$=0.603m) than the motion model ($RMSE_M$=0.923m), ($t(29)=3.74$, $p<0.001$, Cohen's $d=0.49$), very strong evidence for the visual model ($BF_{VM}$ = 40.16). These results confirm the important role of occlusion in explaining pedestrian responses in naturalistic crowd data.
